# Deciphering the metabolic perturbation in hepatic alveolar echinococcosis: a ^1^H NMR-based metabolomics study

**DOI:** 10.1101/320333

**Authors:** Caigui Lin, Lingqiang Zhang, Zhiliang Wei, Kian-Kai Cheng, Guiping Shen, Jiyang Dong, Zhong Chen, Haining Fan

## Abstract

Hepatic alveolar echinococcosis (HAE) is a chronic and potentially lethal parasitic disease. It is caused by growth of *Echinococcus multilocularis* larvae in liver. To date, early-stage diagnosis for the disease is not mature due to its long asymptomatic incubation period. In this study, a proton nuclear magnetic resonance (^1^H NMR) -based metabolomics approach was applied in conjunction with multivariate statistical analysis to investigate the altered metabolic profiles in blood serum and urine samples from HAE patients and to identify characteristic metabolic markers associated with HAE. The current results identified 21 distinctive metabolic difference between the HAE patients and healthy individuals, which can be associated with perturbations in energy metabolism, amino acid metabolism, oxidative stress, and neurotransmitter imbalance. In addition, the Fischer ratio, which is the molar ratio of branched-chain amino acids to aromatic amino acids was found significantly lower (*p*<0.001) in blood serum from HAE patients. The ratio, together with changes in other metabolic pathways may provide new insight into mechanistic understanding of HAE pathogenesis, and may be useful for early-stage HAE diagnosis.

**Author Summary:** Hepatic alveolar echinococcosis (HAE) is a life-threatening disease caused by *Echinococcus multilocularis* infection. The disease has a long asymptomatic early stage (5~15 years), which complicates effective diagnosis of early-stage HAE even with advanced imaging techniques. Metabolomics is an emerging analytical platform that comprises of analysis of all small molecule metabolites that are present within an organism. The applications of metabolomics method on HAE may help to reveal the molecular biology mechanisms of HAE. In the current study, we had used ^1^H NMR-based metabolomics technique to investigate blood serum and urine samples from HAE patients. Altered metabolic responses and characteristic differential metabolites for HAE were identified. The metabolic profiling of human biofluids provided valuable information for early-stage HAE diagnosis and for therapeutic interventions, without having to extract HAE vesicles from patients. By featuring global and comprehensive metabolic status, the metabolomics approach holds considerable promise as a noninvasive, dynamic, and effective tool for probing the underlying mechanism of HAE.

## Introduction

Echinococcosis or hydatid disease is a near-cosmopolitan zoonotic parasitic infection caused by larval stage of pathogenic cestode parasite in the genus *Echinococcus* [1]. Different species of *Echinococcus* cause different diseases. The two main types of diseases are cystic echinococcosis (CE) and alveolar echinococcosis (AE), which are caused by *Echinococcus granulosus* and *Echinococcus multilocularis*, respectively [2]. Compared with CE, AE is associated with less occurrence but higher lethality if timely and proper treatments are unavailable [3]. AE has been reported in definitive hosts (canids) as well as in humans [4].

In humans, AE affects liver in approximately 95% of the diagnosed cases, which is known as Hepatic Alveolar Echinococcosis (HAE) [5]. HAE leads to liver-tissue injury or even hepatic failure primarily through an infiltrative behavior. It resembles the infiltrative proliferation in tumor growth and thus is clinically known as “worm cancer” or “parasite liver cancer”. After a long period of latent and asymptomatic stage, HAE can progress into cirrhotic stage [6] or metastasize into other organs (e.g., lung and brain [7–9]), and cause local organ-function impairment and metastatic infiltration [10].

The HAE disease is endemic in the Northern hemisphere, including North America, Central European and Central Asian countries [11] (e.g. France [12], Germany [13], Austria [14], Poland [15], northern Iran [16], Mongolia [17], and western China [18]). Its global occurrence is mainly attributed to chronic anthropogenic influences, including increased globalisation of animals and animal products, and altered human-animal interfaces [4].

HAE patients often suffer from symptoms including fatigue, abdominal pain, hepatomegaly, nausea and vomiting. Notably, its long asymptomatic period of the early infection stage leads to difficulty in determining time-point or place of infection [19]. In general, HAE diagnosis is based on the integrated considerations of medical history, contact history with suspicious livestock, clinical and pathologic findings, imaging examination, nucleic acid detection and serologic tests [11]. Modern imaging techniques, e.g. ultrasonography (US), magnetic resonance imaging (MRI), fluorodeoxyglucose positron emission tomography (FDG-PET), and computed tomography (CT), have been applied for clinical diagnosis of HAE disease and long-term follow-up after therapeutic interventions [20]. However, these imaging methods have several limitations: CT is unable to delineate disease-induced perihepatic extension; MRI provides good definition of hydatid lesions while fails to assess hydatid fertility (viability); FDG-PET provides noninvasive and accurate evaluation of metabolic activity in HAE at the complexity as well as high expense of administering radiotracer with on-site cyclotron. Besides, effective diagnosis of early-stage HAE remains difficult even with these advanced imaging techniques. This leads to common diagnosis at middle or advanced stage as well as failure to have timely curative resection [21]. Successful early diagnosis and treatment for HAE may prevent complications, reduce postoperative reoccurrence, improve recovery rate, and prove helpful for prognosis.

Metabolomics is an “omics” technique at the downstream of genomics, transcriptomics and proteomics [22]. Among the postgenomic technologies, high-throughput metabolomics has gained increasing attention as it provides accurate determination and quantification of small molecule metabolites in living systems resulting from physiological stimuli or genetic modification [23], which is of diagnostic and prognostic values. Nuclear magnetic resonance (NMR) spectroscopy is one of the leading analytical approaches in metabolomics. It offers unique qualitative and quantitative insight into all of the more abundant compounds present in biological samples [24]. Generally, proton nuclear magnetic resonance (^1^H NMR) -based metabolomics is utilized along with multivariate statistical analysis to capture metabolic states of organisms at the global or “-omics” level and determine characteristic global disease-induced metabolic changes.

In the last decade, metabolomics approaches have been broadly studied and validated in various liver diseases to determine potential early biomarkers and perturbed metabolic pathways [25]. However, reports on applications of ^1^H NMR-based metabolomics method on echinococcosis are limited, except: a ^1^H NMR study to determine metabolites in rodent cyst biomass by Novak *et al*. [26] and an *ex vivo* ^1^H NMR study to explore metabolic profiles with parasite viability in cyst liquid samples from a larger patient collective by Hosch *et al*. [27]. Therefore, a gap of evaluating metabolic variations in biofluids (blood serum and urine) of HAE patients remains to be filled to facilitate the early diagnosis of HAE patients. The metabolic profiling of human biofluids may provide important diagnostic and prognostic information. Blood serum and urine samples serve as two common and important biofluids to be studied in metabolomics due to easy sample collection, pretreatment, restoration, and their ability of giving an overview of global metabolism. In this study, by adopting the high-resolution ^1^H NMR based metabolomics and multivariate statistical analyses, we aim at: firstly, determining biomarkers to differentiate HAE patients from healthy subjects and establishing metabolic fingerprint of biofluids obtained from HAE patients; secondly, consolidating research foundation for molecular biology mechanism of HAE disease; thirdly, demonstrating the feasibility and effectiveness of using metabolomics in echinococcosis studies; finally, providing valuable diagnostic reference for early-stage HAE.

## Results

### ^1^H NMR spectra of biological samples

Representative ^1^H NMR spectra of biofluids collected from the HAE patients and healthy volunteers are shown in Fig 1. In general, several metabolite classes were detected in serum and urine samples, including metabolites involved in energy metabolism, neurotransmission, membrane metabolism, and osmoregulation. In addition, serum metabolome comprised of a few glycoprotein, while a number of gut microbiota-related metabolites were detected in urine samples.

**Fig 1.**
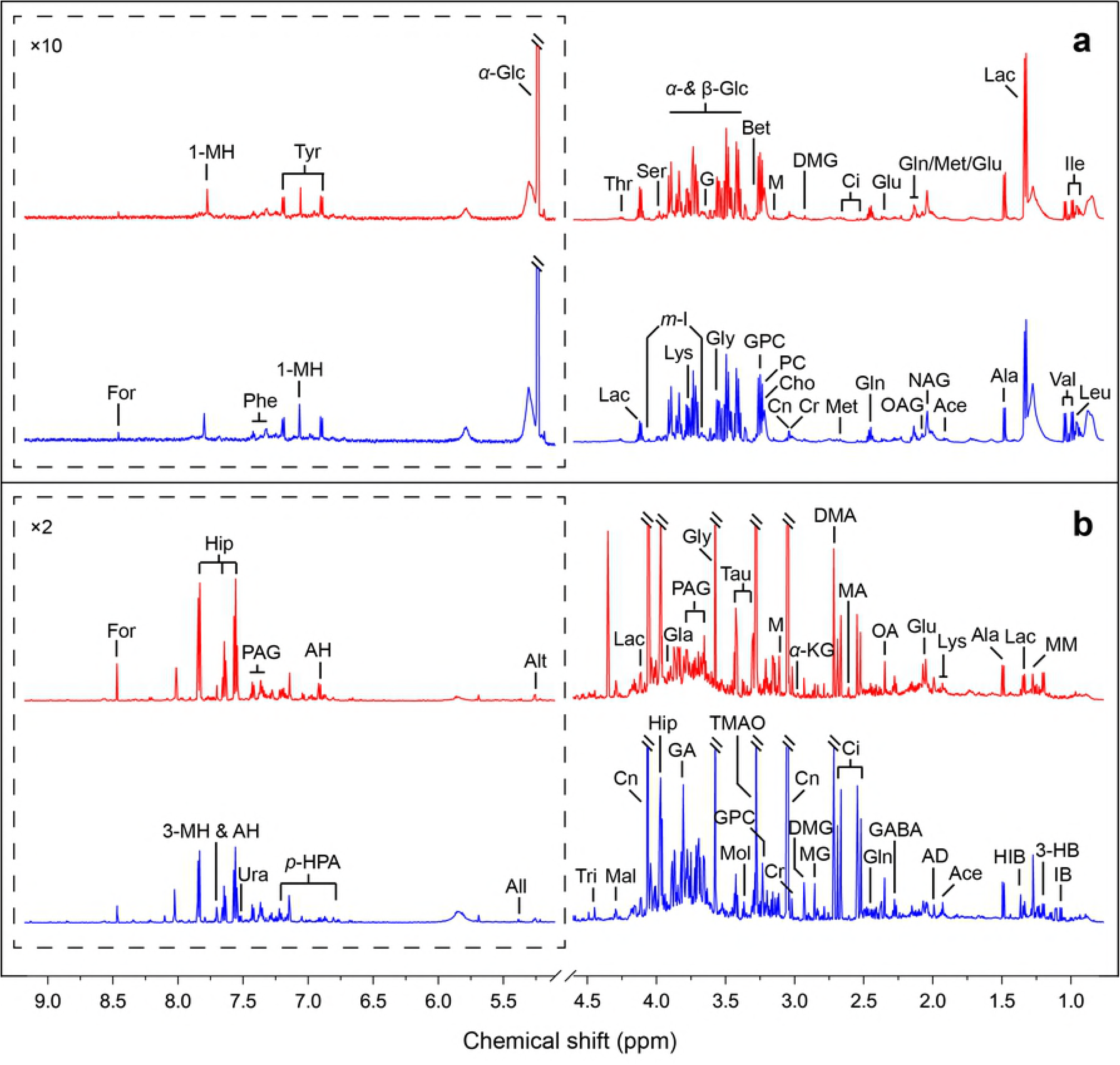
Resonance assignments of representative ^1^H NMR spectra at 600 MHz. (a) serum, (b) urine, HAE (red line) and control (blue line) groups. Spectral regions of 5.2-9.5 ppm for serum and urine samples are displayed with different scales for better visualization of weak and narrow spectral peaks. Assignments: 1-MH, 1-methylhistidine; 3-HB, 3-hydroxybutyrate; 3-MH, 3-methylhistidine; Ace, acetate; AD, acetamide; AH, aminohippurate; Ala, alanine; All, allantoin; Alt, allantoate; Bet, betaine; Cho, choline; Ci, citrate; Cn, creatinine; Cr, creatine; DMA, dimethylamine; DMG, *N,N*-dimethylglycine; For, formate; G, glycerol; GA,; guanidoacetate; GABA, *γ*-aminobutyrate; Gla, glycolate; Gln, glutamine; Glu, glutamate; Gly, glycine; GPC, glycerophosphocholine; HIB, 2-hydroxyisobutyrate; Hip, hippurate; IB, isobutyrate; Ile, isoleucine; Lac, lactate; Leu, leucine; Lys, lysine; M, malonate; MA, methylamine; Mal, malate; Met, methionine; MG, methylguanidine; *m*-I, *myo*-inositol; MM, methylmalonate; Mol, methanol; NAG, *N*-acetylglutamate; OA, oxaloacetate; OAG, *O*-acetylglycoprotein; PAG, phenylacetylglycine; PC, phosphocholine; Phe, phenylalanine; *p*-HPA, *para*-hydroxyphenylacetate; Ser, serine; Tau, taurine; Thr, threonine; TMAO, trimethylamine N-oxide; Tri, trigonelline; Tyr, tyrosine; Ura, uracil; Val, valine; *α*-Glc, *α*-glucose; *α*-KG, *α*-ketoglutarate; *β*-Glc, *β*-glucose.

### Pattern recognition analysis

To uncover HAE-induced metabolic changes, we performed multivariate analyses on the processed NMR data. First, the unsupervised principal components analysis (PCA) was used (Fig 2). Each point in the scores plot in Fig 2 represented an individual serum or urine sample and inter-point distance reflected the scale of their metabolic differences. Moderate groupings could be observed between the control and the HAE groups in the PCA scores plots. By contrast, a distinctive group separation can be achieved when the data was analysed using supervised orthogonal partial least squares discrimination analysis (OPLS-DA). From Fig 3, control group primarily distributed in the left hemisphere, while HAE group fell into the right hemisphere. Such a distinct separation between the control and HAE groups indicated that HAE induced distinctive metabolic changes in human serum and urine. The OPLS-DA models were found robust with excellent predictive powers following permutation tests (200 permutations) (Fig 3).

**Fig 2.**
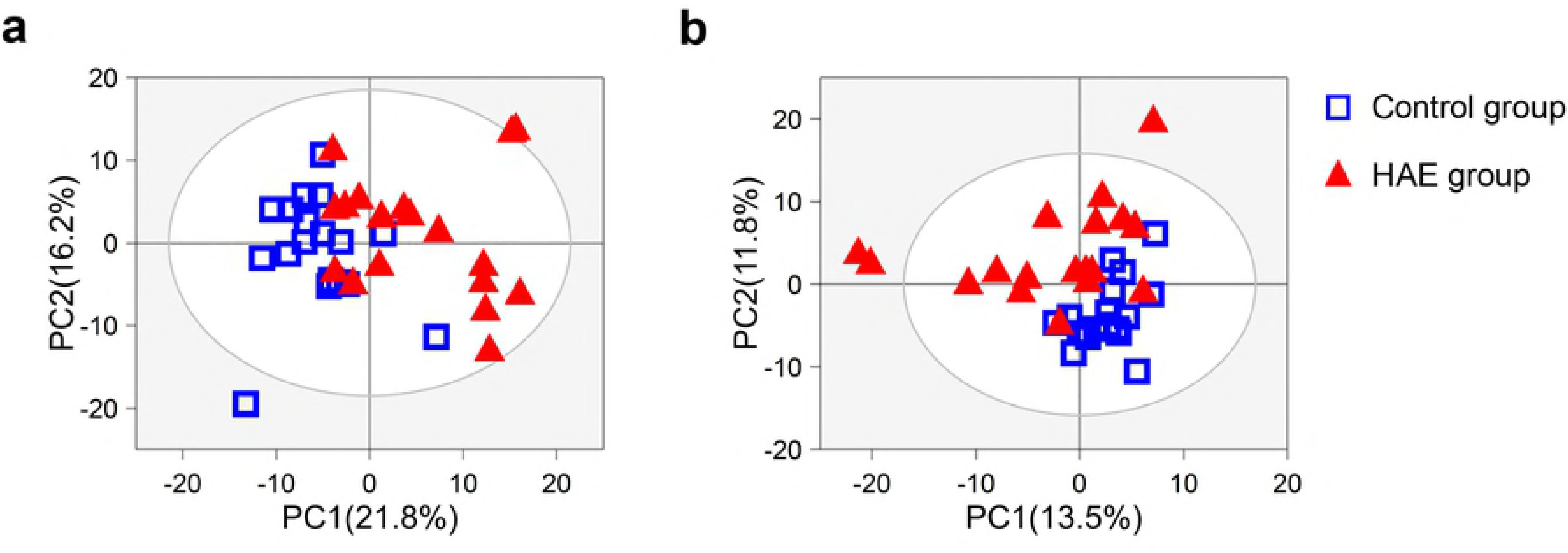
PCA score plots of control and HAE groups. (a) serum and (b) urine. Each data point represents one subject. Two components accounted for 95.5% of total variances in the serum samples and 61.1% of total variances in the urine samples. The outer ellipse represents the 95% confidence interval (Hotelling’s *T*^2^ distribution).

**Fig 3.**
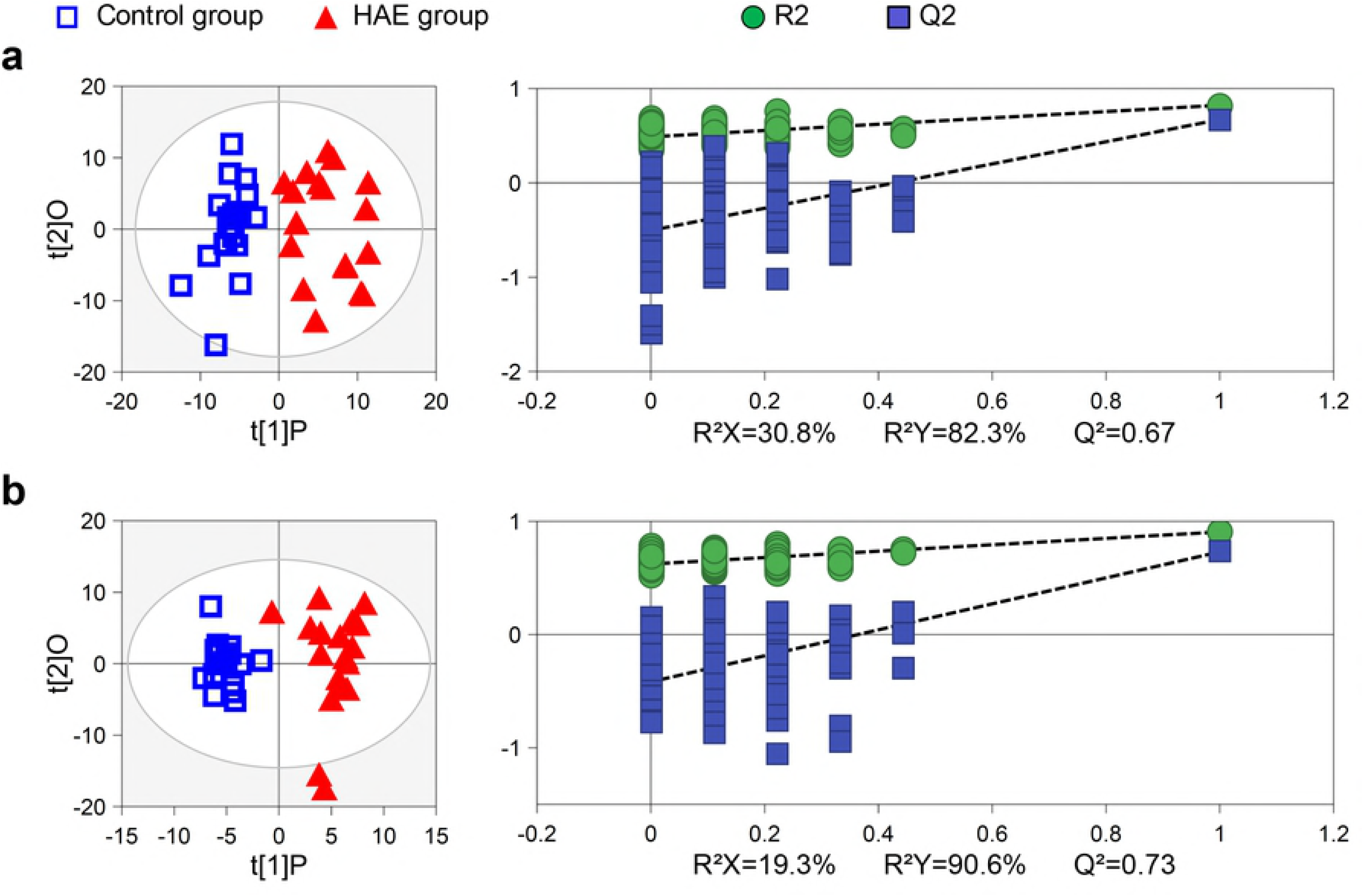
OPLS-DA scores plot (left panel) and the corresponding validation plot (right panel). (a) serum and (b) urine samples. The R^2^ and Q^2^ values reflecting the fraction of variance and the model predictability, respectively, are given to evaluate the model quality.

### Determination of distinctive metabolites for HAE

Next, an enhanced volcano plot was used to identify metabolic markers that differentiate the HAE patients from the healthy controls [28]. In the volcano plot, the importance and significance of the metabolic changes were determined using the following criteria: variable importance projection (VIP) > top 30%, absolute correlation coefficient values (|*r*|) > 0.5, -*log*_10_ (*p*-value) > 2 (i.e., *p* < 0.01), and absolute *log*_2_ (fold change) > 0.25. Generally, identified metabolites with significant changes (those have significant *p*-value, high fold change, VIP, and |*r*|) by multivariate statistical analyses tended to locate at the upper-left or upper-right zones of the enhanced volcano plot in larger circle shapes and warmer colors, as marked with abbreviations in Fig. 4.

**Fig 4.**
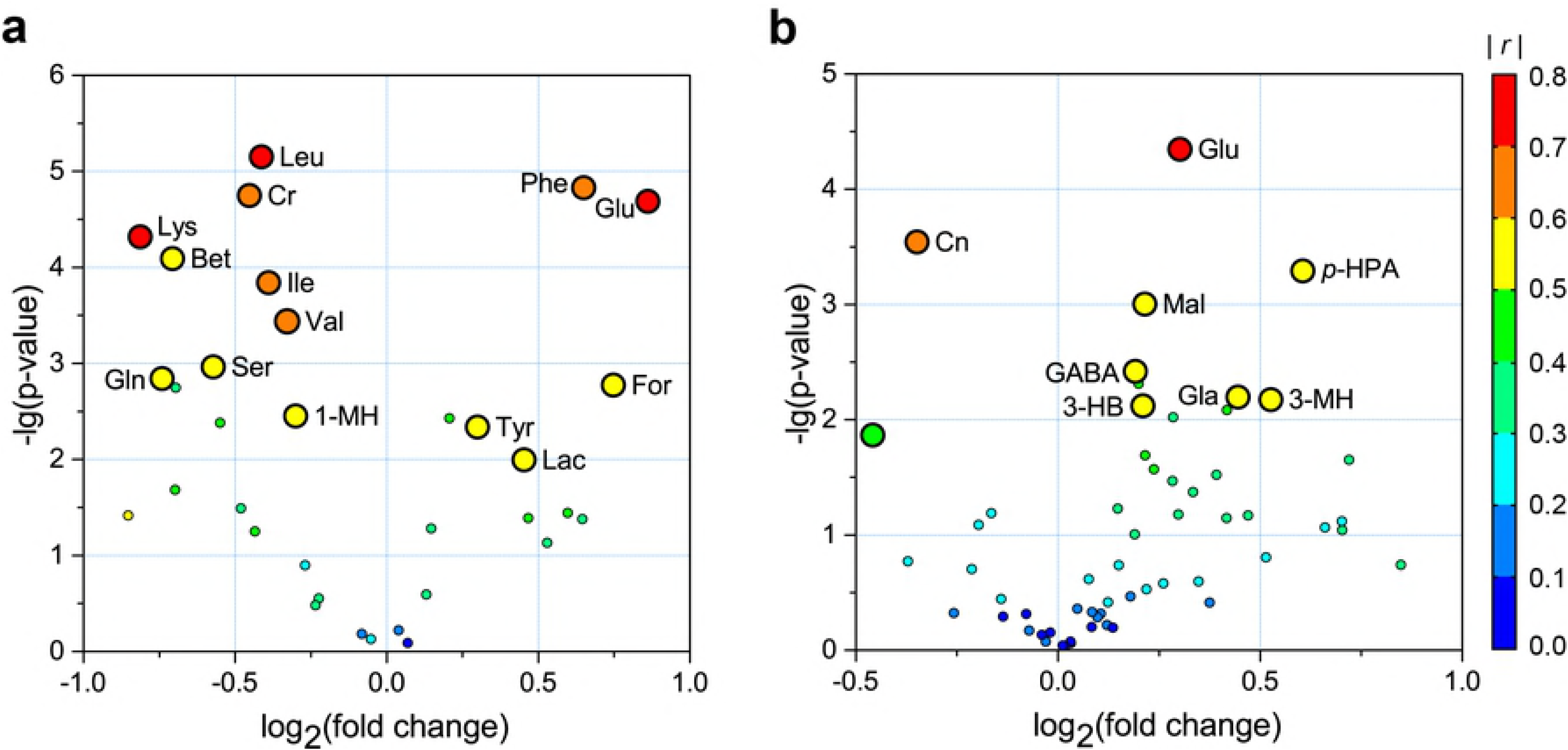
Enhanced volcano plots showing significantly-changed metabolites. (a) serum and (b) urine samples. Volcano plot shows -*log_10_* (*p*-value) on *y*-axis versus *log_2_* (fold change) on *x*-axes. Each point represents a different metabolite. The circles size and color are determined based on the variable importance projection (VIP) and absolute correlation coefficient values (|*r*|), respectively. For each comparison, VIP values are categorized into two categories: top 30% and rest 70 % with each represented by large and small circles, respectively. Warmer color corresponds to higher |*r*|.

For blood serum samples, the enhanced volcano plot showed significant increases of phenylalanine (Phe), glutamate (Glu), tyrosine (Tyr), formate (For), and lactate (Lac), together with decreases of valine (Val), leucine (Leu), isoleucine (Ile), lysine (Lys), serine (Ser), glutamine (Gln), betaine (Bet), creatine (Cr), and 1-methylhistidine (1-MH) in the HAE group (Fig 4a). Using similar approach, urine samples from the HAE group were characterized by significantly higher levels of glutamate (Glu), malate (Mal), glycolate (Gla), 3-hydroxybutyrate (3-HB), *γ*-aminobutyrate (GABA), 3-methylhistidine (3-MH), and *p*-hydroxyphenylacetate (*p*-HPA), as well as lower concentration of creatinine (Cn) (Fig 4b).

### Statistical power analysis

The statistical power analysis of the selected metabolites with significant changes are presented with Table 1. First, the absolute difference between two independent means (the HAE and control groups) and the intra-group standard deviation were used to calculate the effect size. The effect size was considered as large at a value of 0.80 according to the report by Cohen [29]. In addition, post hoc analysis of the achieved statistical power calculation yielded an average value of 0.92 (SD = 0.09) under the given condition of significance level *α* = 0.05, sample size *n* = 18, and specified effective sizes. The reliability and effectiveness of the current results were validated based on sufficiently high effect sizes and statistical power.

**Table 1.**
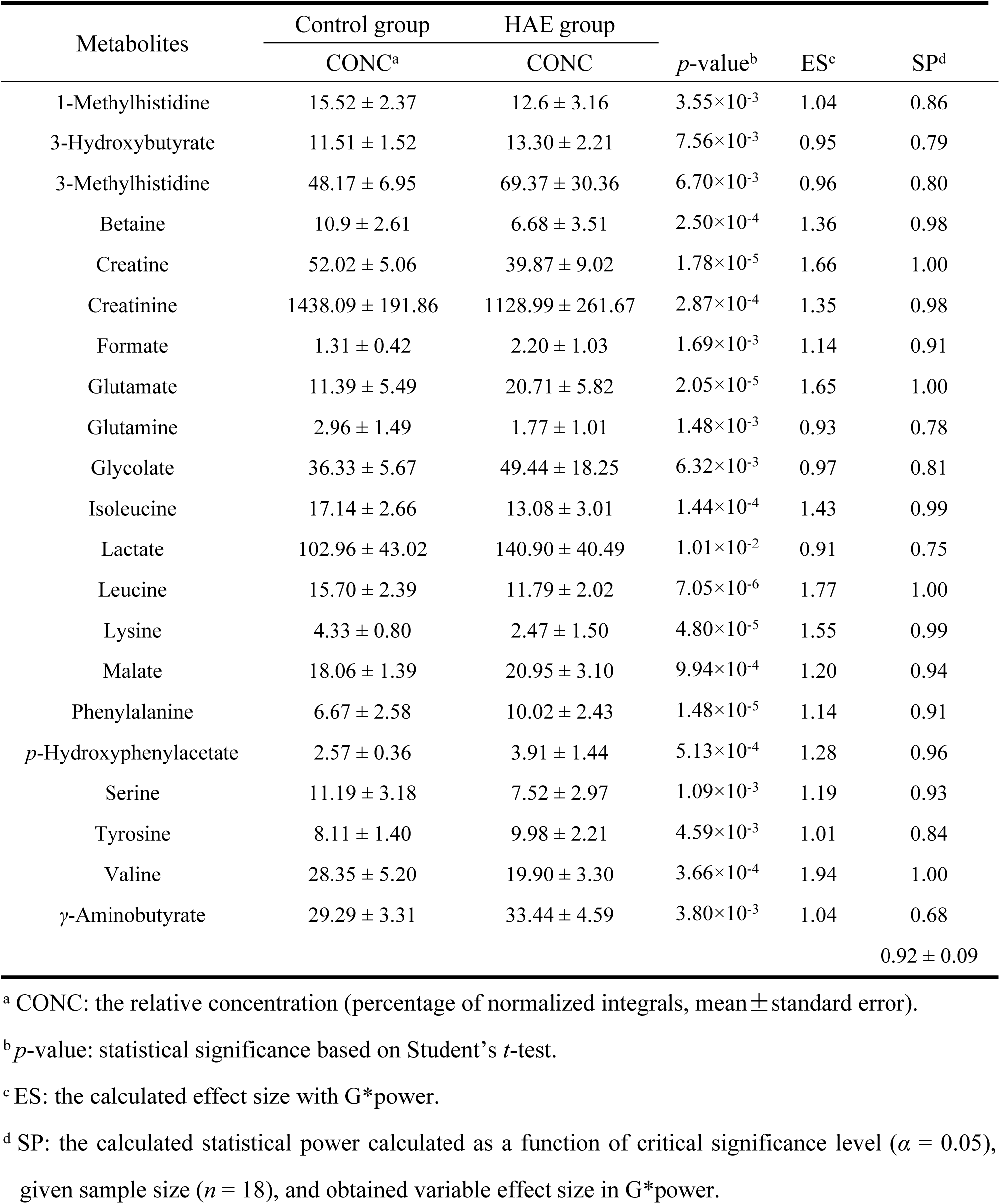
Post-hoc power analyses of characteristic metabolites with G*power.

### Metabolic pathway analysis

A metabolic pathway analysis of differential metabolites between HAE and control groups was performed using the MetaboAnalyst 3.0 software to investigate the most perturbed metabolic pathways in HAE. We found that characteristic metabolites represent perturbations in multiple pathways including alanine, aspartate and glutamate metabolism, arginine and proline metabolism, D-glutamine and D-glutamate metabolism, glycine, serine and threonine metabolism, glyoxylate and dicarboxylate metabolism, lysine metabolism, methane metabolism, phenylalanine metabolism, tyrosine metabolism and valine, leucine and isoleucine metabolism (Fig 5). These pathways were selected based on Impact value ≥ 0.02 and –*log*(*p*) ≥ 5 and considered as most relevant pathways in HAE (Table 2).

**Fig 5.**
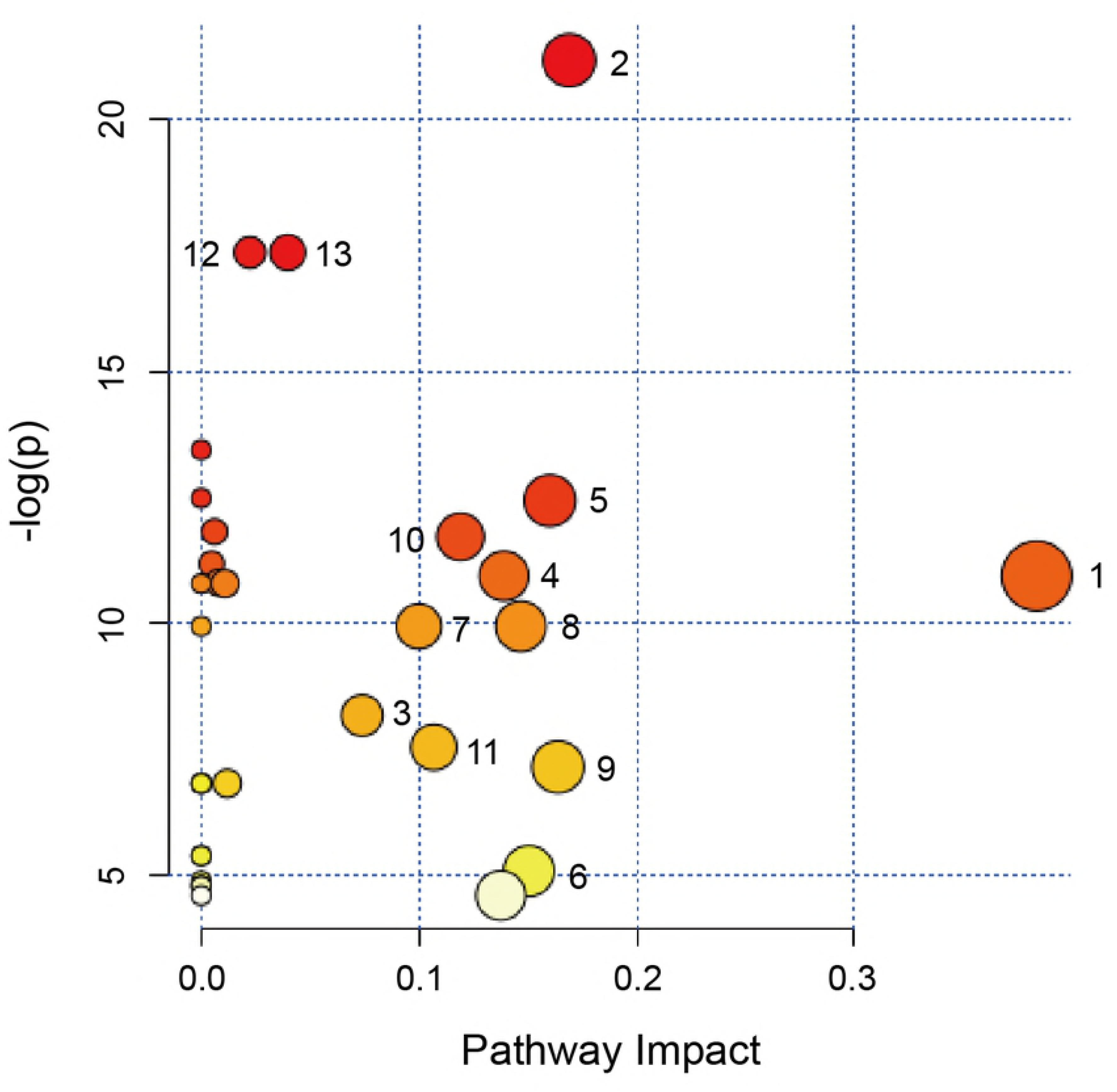
Bubble plots of altered metabolic pathways in bio-samples of HAE compared with the control. Bubble area is proportional to the impact of each pathway with color denoting the significance from highest (in red) to lowest (in white). 1, alanine, aspartate and glutamate metabolism; 2, aminoacyl-tRNA biosynthesis; 3, arginine and proline metabolism; 4, D-glutamine and D-glutamate metabolism; 5, glycine, serine and threonine metabolism; 6, glyoxylate and dicarboxylate metabolism; 7, lysine biosynthesis; 8, lysine degradation; 9, methane metabolism; 10, phenylalanine metabolism; 11, tyrosine metabolism; 12, valine, leucine and isoleucine biosynthesis; 13, valine, leucine and isoleucine degradation.

**Table 2.** Pathway analysis result for metabolomics data with MetaboAnalyst.

| Key | Pathway name | Total<sup>a</sup> | Hit<sup>b</sup> | Impact<sup>c</sup> | $-\log(p)$ |
| --- | --- | --- | --- | --- | --- |
| 1 | Alanine, aspartate and glutamate metabolism | 24 | 2 | 0.38 | 10.94 |
| 2 | Aminoacyl-tRNA biosynthesis | 75 | 9 | 0.17 | 21.18 |
| 3 | Arginine and proline metabolism | 77 | 4 | 0.07 | 8.17 |
| 4 | D-glutamine and D-glutamate metabolism | 11 | 2 | 0.14 | 10.94 |
| 5 | Glycine, serine and threonine metabolism | 48 | 3 | 0.16 | 12.44 |
| 6 | Glyoxylate and dicarboxylate metabolism | 50 | 2 | 0.15 | 5.08 |
| 7 | Lysine biosynthesis | 32 | 1 | 0.10 | 9.94 |
| 8 | Lysine degradation | 47 | 1 | 0.15 | 9.94 |
| 9 | Methane metabolism | 34 | 2 | 0.16 | 7.15 |
| 10 | Phenylalanine metabolism | 45 | 3 | 0.12 | 11.72 |
| 11 | Tyrosine metabolism | 76 | 2 | 0.11 | 7.54 |
| 12 | Valine, leucine and isoleucine biosynthesis | 27 | 3 | 0.04 | 17.37 |
| 13 | Valine, leucine and isoleucine degradation | 40 | 3 | 0.02 | 17.37 |

<sup>a</sup> Total: total number of compounds in the pathway.
<sup>b</sup> Hits: number of the significant different metabolites in the pathway.
<sup>c</sup> Impact: pathway impact value calculated from pathway topology analysis (Betweenness).
<sup>d</sup> $-\log(p)$ : logarithm of the $p$ value calculated by Fisher exact test.

## Discussion

The current study revealed the metabolic perturbation in response to HAE disease by using ^1^H NMR based metabolomics technique. Firstly, ^1^H NMR spectra and pattern recognition analyses of serum and urine showed substantial changes of human metabolome due to HAE. Secondly, a total number of 22 metabolites were identified with moderate ability in discriminating HAE patients from healthy individuals. These metabolites were determined from the enhanced volcano plot, which is the visualization and integration of multivariate and statistical analysis results. Third, metabolic network analyses revealed statistically-significant HAE-induced modulations to a series of metabolic pathways, i.e., amino acid metabolism, energy metabolism, glyoxylate and dicarboxylate metabolism, and methane metabolism (Fig 5 and 6). These metabolic profiles of the host serum and urine opened important perspectives towards understanding biological changes that occur during HAE infection.

**Fig 6.**
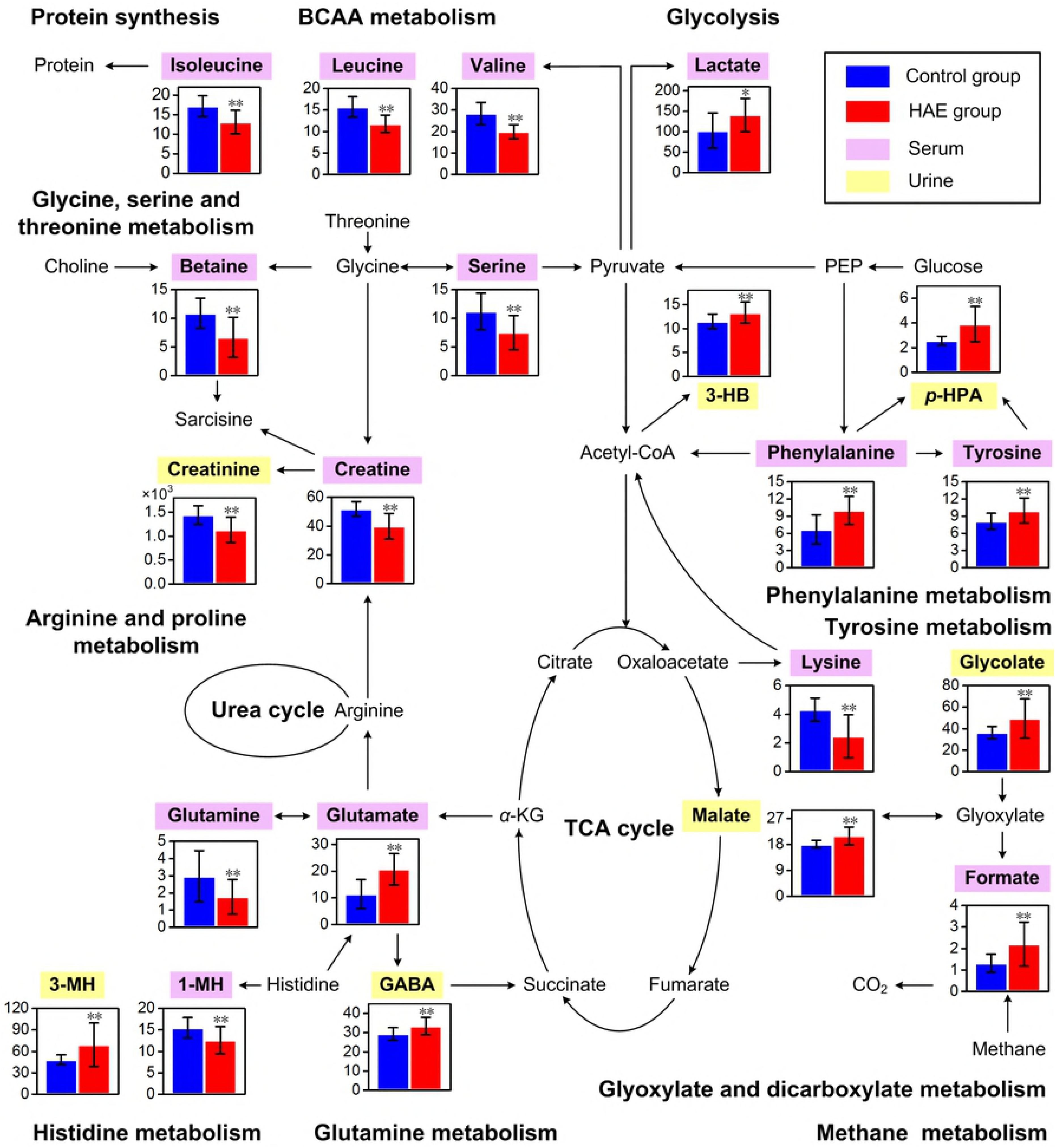
Mapping of significantly-altered metabolites induced by HAE onto plant biosynthetic pathways and potential cross-talks. Each bar graph represents one metabolite in relative concentration (mean±standard error) of control (blue) and HAE (red) groups. * indicates *p* < 0.05 statistical significance relative to control group; ** indicates *p* < 0.01 statistical significance relative to control group.

Alveoli, host metastasis, and massive immune response due to HAE are known to lead to generation of large amount of unstable reactive oxygen species (ROS) [11, 30]. Over-generation of ROS not only affects normal mitochondrial function but also affects cellular homeostasis in case of infection, sterile damage, and metabolic imbalance [31, 32]. This will produce a status of oxidative stress with enhanced energy metabolism and increased protein degradation due to cell damage or necrosis.

This oxidative-stress response often upregulates tricarboxylic acid (TCA) cycle and glycolysis to produce more ATP for cell maintenance and proliferation. The increased levels of malate and lactate in the HAE group than the control group support the notion of enhanced TCA cycle and glycolysis. The decreased creatine concentration in the HAE group provides further support of perturbed energy metabolism in liver mitochondria during HAE infection. Creatine acts as intracellular high-energy phosphate shuttle and plays a crucial role in maintaining cellular energy homeostasis. It is known that a portion of creatine, taken up by cells from the blood through a specific Na^＋^ and Cl^−^ dependent transporter. It can then be phosphorylated to form creatine phosphate, which accepts the high energy phosphate group from ATP and subsequently catabolizes into creatinine to provide an immediate adequate ATP levels in case of energy deficit [33]. Thus, decreases of creatine in blood serum may reflect their consumption to alleviate HAE-induced energy deficit. Apart from being an ergogenic aid, creatine has cytoprotective effect as a direct radical scavenger against ROS [34]. Therefore, the decline of creatine can partly be attributed to the antioxidant reactions during HAE infections. In view of the cellular energy production and antioxidant activity of creatine in living cells, creatine supplementation has been shown to have beneficial effect in prevention and treatment of a number of muscular, neurological, and cardiovascular diseases, including amyotrophic, lateral sclerosis [35], Parkinson’s disease [36], ischemic brain [37], and congestive heart disease [38]. However, the potential effect of creatine supplementation in the HAE patients remained to be explored.

HAE-induced perturbation in amino acids metabolism is indicated by decreased levels of lysine, serine, glutamine, and branch-chain amino acids (BCAA; i.e., valine, leucine, and isoleucine), together with increased levels of glutamate, and aromatic amino acids (AAA; e.g. tyrosine, and phenylalanine) (Figs 4 and 6). Tyrosine, which is the first product of phenylalanine catabolism, accumulates from the hydroxylation of phenylalanine with the aid of a rate-determining enzyme (i.e., phenylalanine hydroxylase) in liver and kidney. Previously, it was reported that directionalities of plasma-concentration shift of these two metabolites are the same and conversion of phenylalanine to tyrosine is an exclusive function of the liver [39, 40]. Thus, the parallel accumulations of AAA (*i.e*. phenylalanine and tyrosine) in blood serum suggested the possible loss of liver function during HAE infection. As essential amino acids, BCAA account for about 20% of our dietary protein and are key regulators of protein syntheses in animals and humans. BCAA disposal is tightly mediated by phosphorylation and dephosphorylation of the branched-chain *α*-keto acid dehydrogenase complex (BCKDC) [41]. The BCKDC is the rate-limiting enzyme in BCAA oxidative catabolism, the possible activation of BCKDC by HAE increases the BCAA catabolism to promote protein synthesis in case of increased protein degradation due to cell damage or necrosis during HAE infection. Moreover, the role of BCAA in immune function has gained growing research interest in the last decade [42]. BCAA are oxidized by immune cells as fuel sources and then incorporated as precursors for the synthesis of new immune cells, effector molecules, and protective molecules [43]. The host immune response induced by HAE may lead to higher consumption of BCAA to maintain immune function.

In addition, HAE was found to down-regulate muscle function during HAE infection. Down-regulated muscle function can be indicated by increased concentrations of GABA and 3-methylhistidine, as GABA has relaxant effects on muscle tone while 3-methylhistidine is an indicator of muscle catabolism [44]. In general, 3-methylhistidine is released during muscle-protein degradation and is not reutilized for muscle-protein synthesis. This suggested that immune response to HAE requires not only redirection of abundant energy but also metabolic resources from other tissues, especially from skeletal muscle. Hence, the consumption of BCAA in promoting protein synthesis may focus on the effect of muscle-protein synthesis. On the other hand, ninety-eight percent of body creatine is found in skeletal muscle and a constant fraction of the body creatine pool is converted each day to creatinine. Hence, it has long been recognized that urinary creatinine excretion constitutes a good reflection of skeletal muscle mass [45]. The decline of urine creatinine reflects a corresponding muscle mass loss in the HAE patients. The symptom of fatigue for HAE patients may attribute to muscle mass loss and down-regulated muscle function.

As large neutral amino acids, BCAA and AAA travel across blood-brain barrier into the brain through a process competing for a same kind of neutral amino acid transporter LAT-1. [46]. Therefore, the raised AAA (tyrosine, and phenylalanine) and reduced BCAA (valine, leucine, and isoleucine) levels in blood serum of the HAE patients may contribute to a subsequent increased influx of tyrosine and phenylalanine in brain during HAE infection. However, the production of important bioamine neurotransmitters (i.e., dopamine, epinephrine and norepinephrine) is relatively insensitive to cerebral levels of their precursor (i.e., tyrosine and phenylalanine). By contrast, the production of false neurotransmitters (e.g., octopamine or phenylethanolamine) will be easier stimulated by the cerebral elevated levels of tyrosine and phenylalanine, leading to increased amount of false neurotransmitters. These false neurotransmitters are structurally similar to “authentic” neurotransmitters, but without ability in propagating neural excitation. This may result in imbalances of neurotransmitter synthesis. Homeostasis between excitatory (e.g., glutamate) and inhibitory (e.g., GABA) neurotransmitters is essential for maintaining the normal functioning of central nervous system (CNS) [47]. In general, glutamate production originates from two pathways: synthesized from *α*-ketoglutarate (one of intermediates of TCA cycle) by the catalysis of glutamate synthase or glutamine-glutamate conversion, and GABA can be produced from glutamate by the glutamate decarboxylase. In our study, elevations of excitatory glutamate and inhibitory GABA along with glutamine reduction indicated that neurotransmitter recycling disorder occurs in the CNS during HAE infection due to the imbalance between excitation and inhibition. Moreover, alterations of GABA and glutamate in the same direction have been reported to contribute to pathophysiology of depression based on behavioral observations [48, 49].

Previously, Fischer *et al*. had reported decreased BCAA and increased AAA levels in blood during hepatic failures in patients with liver disease. The ratio between the three BCAA (valine, leucine, and isoleucine) and two AAA (tyrosine, and phenylalanine) has been termed as Fischer ratio for a quick assessment of liver disease [50]. With a simultaneous consideration over variations of BCAA and AAA, the amino acid imbalance will lead to a lower Fischer ratio in hepatic failures [51–53]. In this study, a decreased Fischer ratio of serum samples for HAE group (*p*＜0.001) indicated that HAE may have triggered hepatic failure in patients. Therefore, Fischer ratio may emerge as a reliable index for diagnosing HAE. Note that a precise Fischer ratio for HAE patients was not available in our experiments, but this does not affect the effectiveness of Fischer ratio in discriminating HAE patients from healthy individuals. Moreover, the inadequate development of HAE among patients and high altitude subjects live in our study may partly attribute to the difference in Fischer ratio as compare with results of other researchers.

Based on these analyses, the imbalanced BCAA and AAA in serum play a crucial pathophysiologic role in HAE. A raise in serum levels of BCAA is anticipated to restore a normal Fischer ratio and reverse the imbalance, aiding the HAE patients in promoting protein synthesis, improving muscle function, protecting against stress, maintaining neurotransmitter equilibrium and strengthening immune response.

It should be noted that the present study has several limitations. First, a larger number of subjects will further increase the reliability and accuracy of current results. Then, an integrated application of NMR with LC-MS or GC-MS will extend the metabolite coverage, and multiple -omic techniques (e.g., genomics and proteomics) can cross-validate and better interpret the experimental results. Finally, a distinctive method that can distinguish HAE from other types of hepatic disease is not available with the current study. Such a method will prove helpful in avoiding clinic misdiagnoses since HAE exhibits certain similarities (e.g., lower Fischer ratio) with other hepatic disease. The possibility of using Fischer ratio to categorize different types of hepatic diseases and unique metabolic marker for HAE apart from other hepatic diseases will be explored in our future study, where groups with non-HAE hepatic diseases will be included.

The attractive properties of NMR based metabolomics approach, i.e., excellent repeatability, short detection duration, and multiple metabolite coverage in a single measurement, facilitate the HAE studies. A systematic exploration over multiple metabolites in small molecules helps to reveal the comprehensive metabolic variations induced by HAE infection. The early-stage HAE diagnose may be possible with a combination of both metabolomics and imaging techniques. For instance, chemical-exchange weighted magnetic resonance imaging techniques can be utilized to yield maps weighted by metabolite of interest [54–56]. Such maps are obtained through an exchange effect with water and thus are characterized by significant signal enhancement to observe small metabolic variations unnoticeable with other imaging modalities. A general issue in such techniques is the lack of specificity, and modulations from adjacent resonances are inevitably and unfavorably introduced. The NMR based metabolomics with high specificity can thus be combined for analyses and better interpret the result by mutual reference.

In conclusion, ^1^H NMR-based metabolomics approach was used to investigate endogenous metabolic characterizations of HAE disease, thus helping to reveal the molecular biology mechanisms of HAE. Particularly, multivariate statistical analyses have highlighted several characteristic metabolic makers for HAE, e.g., decreased BCAA (valine, leucine, and isoleucine) group, increased AAA (tyrosine, and phenylalanine) group, increased lactate, and decreased creatine to list a few. The altered Fischer ratio holds high diagnostic potential as a predictor for evaluating the status or future development of HAE. These findings may provide valuable reference information for early-stage HAE diagnoses and for therapeutic interventions. By featuring global and comprehensive metabolic status. The current metabolomics approach holds considerable promise as a noninvasive, dynamic, and effective tool for probing the underling mechanism of HAE.

## Materials and Methods

### Ethical statement

All procedures involving human subjects were approved by the ethical committee of Affiliated Hospital of Qinghai University in Xining, Qinghai, China. Written informed consents were obtained from all subjects involved in the study.

### Participant recruitment

The recruited HAE patients were diagnosed at the Hepatopancreatobiliary Surgery Department in the Affiliated Hospital of Qinghai University from July to September 2016. Diagnoses of HAE relied on integrated usage of imaging examination, serologic test, nucleic acid detection and pathologic observation. In addition, patients’ medical history and contact with potential animal host of the disease were also taken into consideration. Healthy volunteers from patients’ families were recruited as the control group. To prevent confounding metabolic effect from other diseases, patients with the following diseases were excluded from the study: diabetes, nephrosis, autoimmunity disease, malignant hepatic tumor, fever, severe hepatorenal dysfunction, and post-transplantation immunoreaction.

Sample size used in this study was estimated based on the prior power test with some knowledge from our previous preliminary study. The power analyses was done using MetaboAnalyst 3.0 software (http://www.metaboanalyst.ca) [57, 58], and the result indicated that sample size of 18 is adequate to provide sufficiently high statistical power (Fig 7).

**Fig 7.**
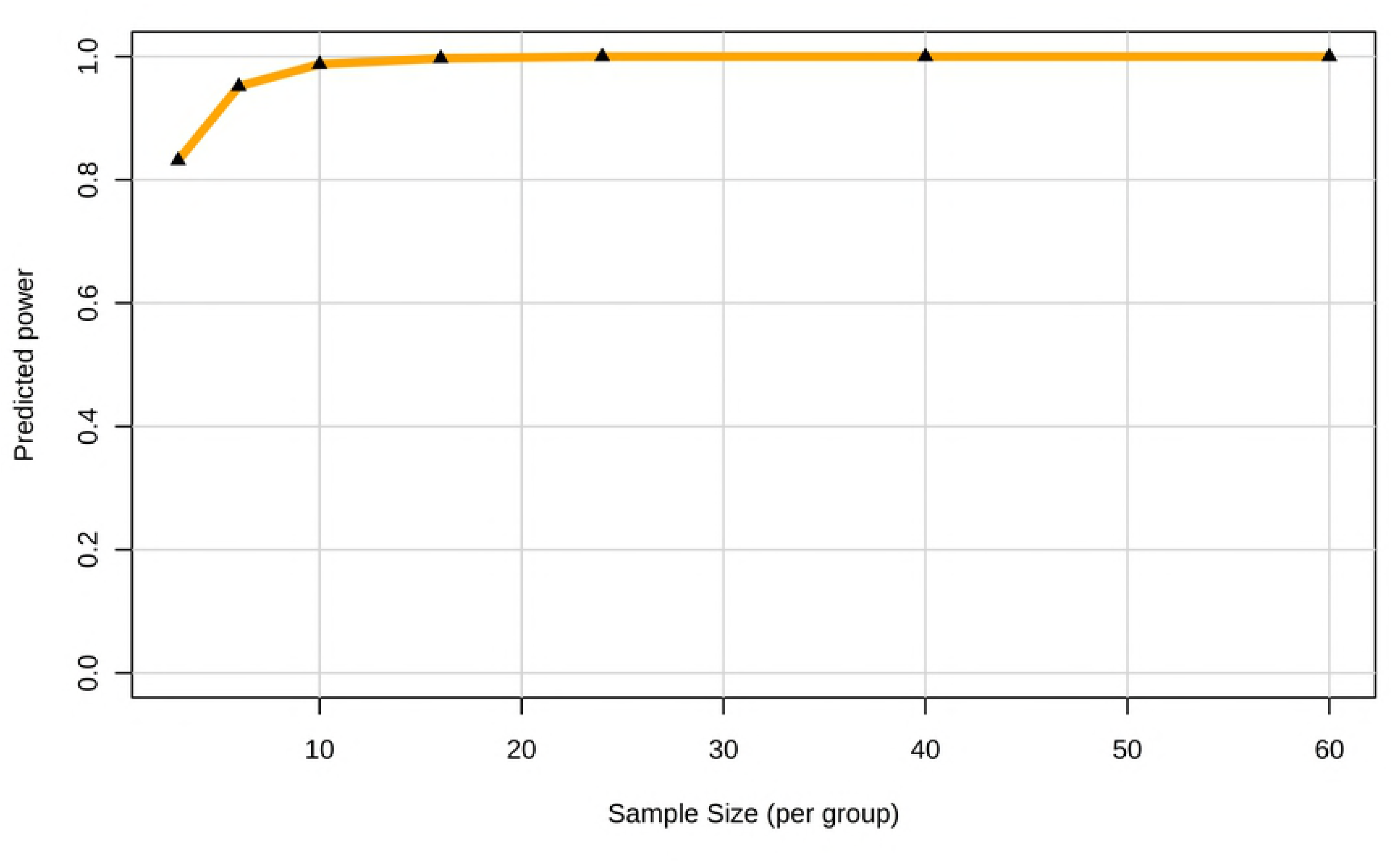
Sample size profile with false discovery rate (FDR)=0.005.

### Sample collections and preparations

All blood and urine samples were collected in the morning, 12 hours after the last meal of the previous day (fasting conditions). Blood samples were collected into tubes without anticoagulant and left to clot at room temperature for 20 minutes. Then blood samples were centrifuged at 11000×g at 4 °C for 15 minutes to obtain blood serum. Then, the blood serum and urine samples were aliquoted, snap-frozen in liquid nitrogen and stored at −80 °C until further treatments.

Prior to analysis, samples were thawed at room temperature. An aliquot of 400 μL blood serum was added with 200 μL phosphate buffer solution (90 mM K_2_HPO_4_/NaH_2_PO_4_, pH 7.4, 0.9% NaCl, 99.9% D_2_O). On the other hand, 300 μL of urine samples were mixed with a different phosphate buffer solution (300 μL, 1.5 M K_2_HPO_4_/NaH_2_PO_4_, pH 7.4, 99.9% D_2_O with 0.3 mM 3-trimethylsilyl-propionic-2,2,3,3-d4 acid (TSP) as a chemical-shift reference for 0 ppm). D_2_O was used to provide the NMR spectrometer with a field frequency for locking. Buffered serum and urine samples were then centrifuged (11000×g, 4 °C, 10 min) to remove debris, and 500 μL supernatant from each 600 μL mixture was transferred into 5-mm NMR tube. In total, 36 serum and urine samples in NMR tubes were prepared and stored at 4 °C before NMR analysis.

### ^1^H NMR experiment

^1^H NMR experiments were performed using a Bruker NMR system operating at the proton frequency of 600 MHz. The operating temperature was set at 298K. The Carr-Purcell-Meiboom-Gill (CPMG) sequence (awaiting time ~ π/2 ~ [τ ~ π ~τ]_n_ ~ acquisition) was used to acquire spectra of blood serum samples with an echo time (τ) of 250 μs and a free relaxation duration (2nτ) of 100 ms. For urinary samples, Nuclear Overhauser Effect Spectroscopy (NOESY, awaiting time ~ π/2 ~ t_1_ ~ π/2 ~ t_m_ ~ π/2 ~ acquisition) was implemented with a 2 s water suppression and mixing time (t_m_) of 120 ms. Free induction decays were recorded into 32 k data points with 64 averages at a spectral width of 10 kHz prior to Fourier transformation.

### Data processing of ^1^H NMR spectra

Data pre-processing for the acquired ^1^H NMR spectra (including Fourier transformation, baseline correction, and phase correction) was performed using the MestReNova v.8.1.2 software (Mestrelab Research S.L.). TSP signal was set as δ 0.00 for urine sample, and left split of the doublet of lactate signals was set as δ 1.336 for serum samples. Residual water signals (serum: δ 4.65-5.15, urine: δ 4.75-5.15), urea resonances (δ 5.70-6.40), and peak-free regions were selectively excluded from further analyses. The remaining spectra over ranges of δ 0.8-8.5 for blood serum and δ 0.8-9.5 for urine were segmented into bucketed data using self-adaptive integration [59], and the results were saved in Microsoft Excel files. For data normalization, the probabilistic quotient normalization (PQN) [23] was utilized to compensate for overall concentration variations. The peaks in the acquired ^1^H NMR spectra were assigned based on previous published literatures [60, 61], the KEGG database [62], and the HMDB database [63]. The relative concentrations for all assigned metabolites were evaluated based on their normalized peak integral area. For metabolites that gave rise to multiple peaks, peaks in the least overlapping spectral region were chosen for quantification.

### Multivariate and univariate statistical data analyses

Multivariate analyses of the pre-processed data were performed using the SIMCA software (version 14.1, Umetrics, Umeå Sweden). The data were examined by the non-supervised principal components analysis (PCA, unit variance scaling) and orthogonal partial least squares-discrimination analysis (OPLS-DA, unit variance scaling). A 7-fold cross-validation and permutation test (200 permutations) were performed and the obtained values of R^2^ (total explained variation) and Q^2^ (model predictability) were applied to validate the constructed models [64]. In general, model with Q^2^ ≥ 0.4 is considered as a reliable model [65]. The variable importance projection (VIP) and absolute correlation coefficient (|*r*|) constructed from the OPLS-DA analysis were used as parameters to select differential metabolites. VIP, which was denoted as a unitless number, delineates the contribution of each predictor variable to the model and presents the influence of each predictor on response variables. A larger VIP corresponds to a greater discriminatory power for the metabolites. Student’s *t*-test constitutes a simple statistical analyses to determine statistically significant metabolic variations in univariate analysis with transformed *p*-value [66]. For a metabolite, its fold change was calculated based on the ratio of average concentrations between the HAE group and the control group.

In this study, results of multivariate statistical analysis were visualized with volcano plot to identify metabolites with significant difference between the HAE and control groups [67, 68]. Particularly, the interactive volcano plot with vertical axis denoting -*log_10_* (*p*-value) and horizontal axis denoting *log_2_* (fold change) presents circles with different size and color for displaying VIP and `|r|` values, respectively.

Additionally, a post hoc power analysis with specified significance level *α*, sample size, and effect size was executed with the online statistics software G*power 3.1 (http://www.gpower.hhu.de/) [69, 70] to demonstrate the statistical power of reported results.

## Acknowledgments

We thank all the members for helpful discussions.

## Supporting Information Legends

**File 1. (Serum.xlsx): Binned ^1^H NMR spectra of serum samples.**

**File 2. (Urine.xlsx): Binned ^1^H NMR spectra of urine samples.**

